# Metabolic Signatures of Performance in Elite World Tour Professional Cyclists

**DOI:** 10.1101/2022.09.13.507793

**Authors:** Travis Nemkov, Francesca Cendali, Davide Stefanoni, Janel L. Martinez, Kirk C. Hansen, Iñigo San-Millán, Angelo D’Alessandro

**Affiliations:** University of Colorado Anschutz Medical Campus, Department of Biochemistry and Molecular Genetics, Aurora, CO, USA; University of Colorado Anschutz Medical Campus, Department of Medicine, Division of Endocrinology, Metabolism and Diabetes. Aurora, CO, USA; University of Colorado, Colorado Springs, Department of Human Physiology and Nutrition, Colorado Springs, CO, USA

**Keywords:** exercise, cycling, bioenergetics, metabolomics, lipidomics, glycolysis, fatty acid oxidation, carnitine

## Abstract

**Introduction:** Metabolomics studies of recreational and elite athletes have been so far limited to venipuncture-dependent blood sample collection in the setting of controlled training and medical facilities. However, limited to no information is currently available if findings in laboratory settings are translatable to real world scenario in elite competitions.

**Methods:** To characterize molecular profiles of exertion in elite athletes during cycling, we performed metabolomics analyses on blood isolated from twenty-eight international-level elite World Tour professional male athletes from a Union Cycliste Internationale (UCI) World Team taken before and after a graded exercise test (GXT) to volitional exhaustion and before and after a long aerobic training session. Moreover, established signatures were then used to characterize the metabolic physiology of five of these cyclists that were selected to represent the same UCI World Team during a 7-stage elite World Tour race.

**Results:** Using dried blood spot collection to circumvent logistical hurdles associated with field sampling, these studies defined metabolite signatures and fold change ranges of anaerobic or aerobic exertion in elite cyclists, respectively. Blood signatures derived in controlled settings enabled comparison with blood sampled during competition, thus providing insight into fatigue status of the cyclists during the course of the race. Collectively, these studies provide a unique view of alterations in the blood metabolome of elite athletes during competition and at the peak of their performance capabilities.

**Graphical abstract:** 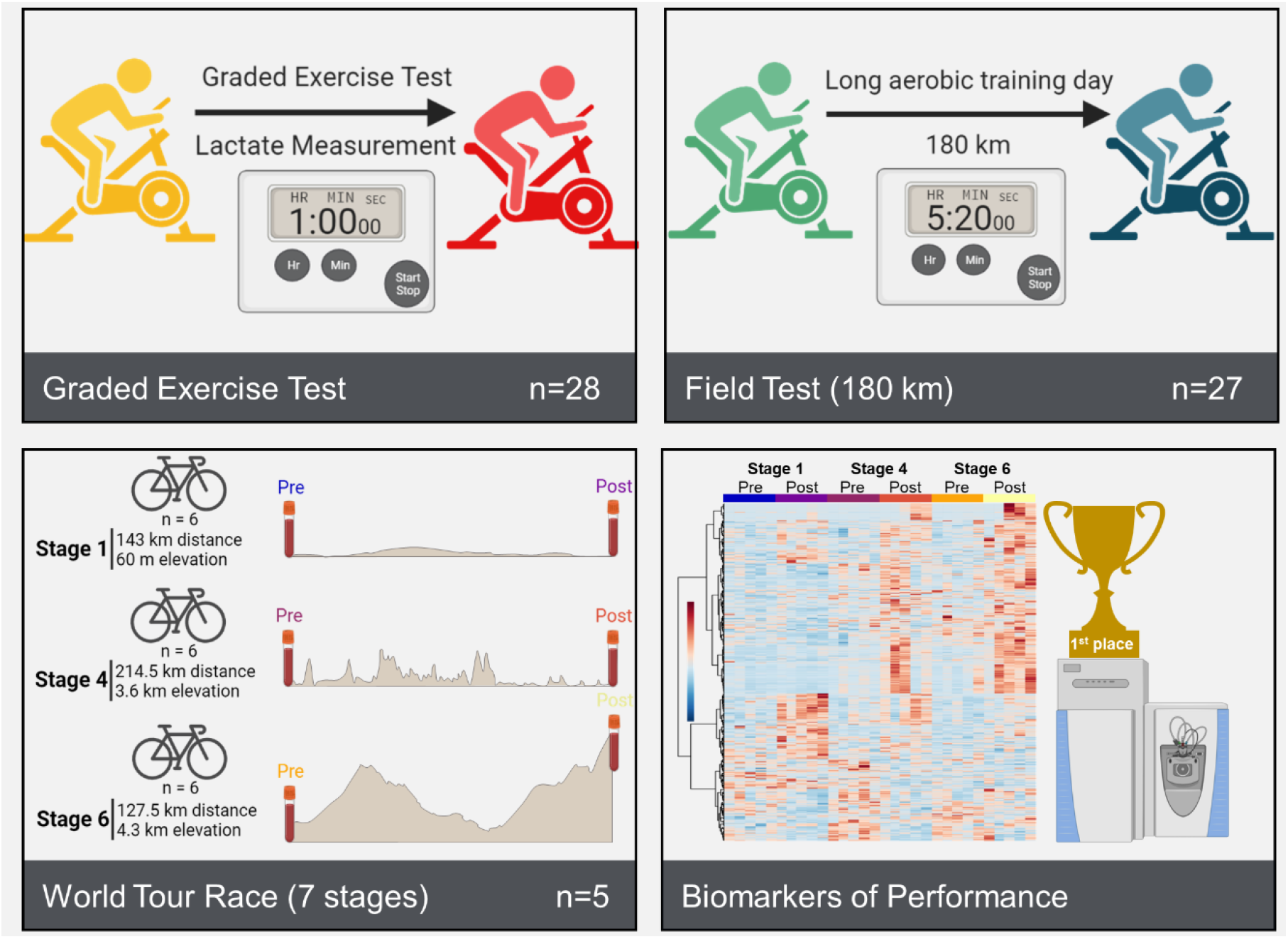

**Summary:** Nemkov et al. leveraged field sampling and blood metabolomics and lipidomics approaches to follow a professional Team of elite cyclists upon graded exercise test to volitional exhaustion, field aerobic training (180 km) and a during multi-stage World Tour race. They identify markers of cycling performance beyond lactate thresholds (ranging from 3.75 to 6.5 watts per kilogram in this group), including carboxylic acids, fatty acids and acylcarnitines.

**Key points:**

- We profiled metabolism of 28 international-level elite World Tour professional male athletes from a Union Cycliste Internationale UCI World Team during training and World Tour multi-stage race;
- Dried blood spot sampling affords metabolomics analyses to monitor exercise performance;
- Determination of lactate thresholds during graded exercise test (GXT) to volitional exhaustion shows a range of from 3.75 to 6.5 watts per kilogram in this group;
- Blood profiles of lactate, carboxylic acids, fatty acids and acylcarnitines differed between different exercise modes (GXT and 180 km aerobic training session);
- Metabolic profiles were affected by stage-specific challenges (sprint vs climbing) during a World Tour multi-stage race.

## 1. INTRODUCTION

Investigations into the metabolic effects of exercise [1,2] have helped elucidate physiological adaptations to stress and have been critical for understanding pathologies in which metabolism is dysregulated. From aging[3] to cardiovascular and other non-communicable diseases[4,5], from cancer[6] to hemorrhagic or ischemic hypoxia[7], from immunometabolism[8] to neurodegenerative diseases[9], metabolic derangements are increasingly appreciated as etiological contributors to disease onset, severity, and prognosis.

Energy requirements and substrate utilization during physical exertion are dependent upon workload and the perfusion of oxygen. During low and medium intensity exercise when oxygen supply is sufficient to meet bioenergetic demands, fatty acid substrates are relied upon for generation of adenosine triphosphate (ATP) through the use of fatty acid oxidation to fuel the Tricarboxylic Acid (TCA) cycle. As exertional intensity increases to a point where oxidative phosphorylation is no longer sufficient, metabolism shifts to favor the rapidity of carbohydrate-driven glycolysis for ATP generation, a metabolic switch that increases the rate of lactate formation. Under high glycolytic rates, lactate production exceeds the rate of lactate consumption in the mitochondria and thus begins to accumulate in circulation [10]. While lactate is primarily formed in fast-twitch fibers, it is oxidized during exercise as a substrate to fuel mitochondria of adjacent slow-twitch muscle fibers. A metabolic inflection point, called the “lactate threshold” (LT), is defined by the maximal effort or intensity that an athlete can maintain for an extended period of time with little or no increase in circulating lactate [11]. As such, classic studies have focused on LT as a proxy for exercise performance.

In the last decade, there has been an increased focus on the metabolic responses to exercise including measurement of substrate utilization, metabolic flexibility and mitochondrial function in athletes[12]. Recent advances in metabolomics have fostered a more comprehensive understanding of human responses to low, moderate or high intensity exercise[13,14]. Results have evidenced a differential impact of exercise modes (e.g., acute versus resistance and endurance exercise[15] and across different sports (e.g., endurance athletes, sprinters, bodybuilders[16,17] on the extent and magnitude of metabolic reprogramming. While we have recently reported metabolomics-based characterizations of cycling in recreational[18] and professional[19] athletes under controlled acute training regimens, literature is scarce on elite professional athletes undergoing testing in the field and, more importantly, during World Tour competitions. Performing such investigations in elite professional athletes offers a unique opportunity to determine the ceiling of human performance, against which we can scale human physiology of healthy occasional, recreational, semi-professional and professional athletes. Such scale would also inform a model on how human metabolism works at its optimum, which in turn could drive the interpretation of metabolic derangements under pathological conditions, even at their early onset.

## 2. METHODS

### 2.1 Ethical statement

All study procedures were conducted in accordance with the Declaration of Helsinki and in accordance with a predefined protocol that was approved by the Colorado Multiple Institutional Review Board (COMIRB 17-1281). Written informed consent was obtained from all subjects.

### 2.2 Graded Exercise Test

Twenty-eight international-level elite World Tour professional male cyclists performed a graded exercise test (GXT) to exhaustion on an electrically controlled resistance leg cycle ergometer (Elite, Suito, Italy). After a 15-minute warm-up, participants started leg cycling at a low intensity of 2.0 W·kg^-1^ of body weight. Exercise intensity was increased 0.5 W·kg^-1^ every 10 minutes as previously described (San-Millán et al., 2009) Power output, heart rate and lactate were measured throughout the entire test and recorded every 10 minutes including at the end of the test.

### 2.3 Blood Lactate concentration measurement

At the end of every intensity stage throughout the graded training period, a sample of capillary blood was collected to analyze both intra- and extra-cellular levels of L-lactate (Lactate Plus, Nova Biomedical, Waltham, MA, USA). Heart rate was monitored during the whole test with a heart monitor (Polar S725x, Polar Electro, Kempele, Finland).

### 2.4 Aerobic Training Session

Two days after the GXT, 27 international-level World Tour professional male cyclists (Tour de France Level) completed a 180 km training ride over 5:20h. During the ride, each cyclist performed 3 climbs at lactate threshold (as determined by the previous maximal physiology test) and then completed the final 40 km at low aerobic intensity.

### 2.5 Blood Sampling for Metabolomics

Twenty µl of whole blood was sampled before and after the Maximal Physiology Test and the aerobic training session using a Mitra® Volumetric Absorptive Sampling (VAMS) device and dried under ambient conditions according to manufacturer’s instructions (Neoteryx, Torrance, CA, USA). Samples were individually sealed in air-tight packaging in the presence of a desiccant and shipped under ambient conditions to the University of Colorado Anschutz Medical Campus. Upon arrival, individual samples were added to 200 µl of methanol:acetonitrile:water (5:3:2 v/v/v) and sonicated for 1 hour as described [20,43]. Metabolite extracts were isolated by centrifugation at 18,000 x g for 10 minutes. Supernatants were separated into autosampler vials.

### 2.6 World Tour Stage Race

Five international-level World Tour professional male cyclists were selected to represent a Union Cycliste Internationale (UCI) World Team and participated to a 7-stage elite World Tour race. Performance parameters such as speed and power output were monitored using Training Peaks (Louisville, CO, USA). Due to logistical constraints, whole blood samples were collected using the TAP device (Seventh Sense Biosystems, Medford, MA) as previously described (Catala et al., 2018) prior to and immediately upon completion of Stages 1, 4, and 6. Samples were frozen in dry ice within 15 minutes of isolation and were maintained under this condition until all samples were collected, upon which they were shipped on dry ice and stored at −80°C until analysis.

### 2.7 Sample Preparation

Prior to LC-MS analysis, samples were placed on ice and re-suspended with 9 volumes of ice cold methanol:acetonitrile:water (5:3:2, v:v) and vortexed continuously for 30 min at 4°C for metabolomics or 9 volumes of ice cold methanol and incubated for 30 minutes at −20°C for lipidomics. Prior to sample resuspension, the metabolomics extraction solution was supplemented with final concentrations of 0.5 µM ^13^C^15^N amino acid mixture, of 40 µM ^13^C_1_-lactate, and 50 µM ^13^C_6_-glucose (Cambridge Isotope Laboratories, Inc., Tewksbury, MA) for determination of absolute molar concentration of analytes. Insoluble material was removed by centrifugation at 18,000 g for 10 min at 4°C and supernatants were isolated for omics analysis by UHPLC-MS. Metabolite extracts were dried down under speed vacuum and re-suspended in an equal volume of 0.1% formic acid prior to analysis, while lipidomics extracts were analyzed in methanol.

### 2.8 UHPLC-MS data acquisition and processing

Analyses were performed as previously published (Nemkov et al., 2017; Reisz et al., 2019). Briefly, the analytical platform employs a Vanquish UHPLC system (Thermo Fisher Scientific, San Jose, CA, USA) coupled online to a Q Exactive mass spectrometer (Thermo Fisher Scientific, San Jose, CA, USA). Polar extracts (2 µl injections) were resolved over a Kinetex C18 column, 2.1 × 150 mm, 1.7 µm particle size (Phenomenex, Torrance, CA, USA) equipped with a guard column (SecurityGuard™ Ultracartridge – UHPLC C18 for 2.1 mm ID Columns – AJO-8782 – Phenomenex, Torrance, CA, USA) using an aqueous phase (A) of water and 0.1% formic acid and a mobile phase (B) of acetonitrile and 0.1% formic acid for positive ion polarity mode, and an aqueous phase (A) of water:acetonitrile (95:5) with 1 mM ammonium acetate and a mobile phase (B) of acetonitrile:water (95:5) with 1 mM ammonium acetate for negative ion polarity mode. The Q Exactive mass spectrometer (Thermo Fisher Scientific, San Jose, CA, USA) was operated independently in positive or negative ion mode, scanning in either Full MS mode (2 μscans) from 60 to 900 m/z at 70,000 resolution, with 4 kV spray voltage, 45 sheath gas, 15 auxiliary gas, AGC target = 3e6, maximum IT = 200 ms or data-dependent fragmentation (Top 15 ddMS2, for validation) at 17,500 resolution, AGC target = 1e5, maximum IT = 200 ms, isolation window = 4.0 m/z, NCE = 20, 24, 28. Non-polar lipid extracts were resolved over an ACQUITY HSS T3 column (2.1 × 150 mm, 1.8 µm particle size (Waters, MA, USA) using an aqueous phase (A) of 25% acetonitrile and 5 mM ammonium acetate and a mobile phase (B) of 90% isopropanol, 10% acetonitrile and 5 mM ammonium acetate. The column was equilibrated at 30% B, and upon injection of 10 μl of extract, samples were eluted from the column using the solvent gradient: 0-9 min 30-100% B at 0.325 mL/min; hold at 100% B for 3 min at 0.3 mL/min, and then decrease to 30% over 0.5 min at 0.4 ml/min, followed by a re-equilibration hold at 30% B for 2.5 minutes at 0.4 ml/min. The Q Exactive mass spectrometer (Thermo Fisher) was operated in positive ion mode, scanning in Full MS mode (2 μscans) from 150 to 1500 m/z at 70,000 resolution, with 4 kV spray voltage, 45 sheath gas, 15 auxiliary gas, AGC target = 3e6, maximum IT = 200 ms. Data-dependent fragmentation (Top 10 ddMS2) was performed at 17,500 resolution, AGC target = 1e5, maximum IT = 50 ms, and stepped NCE of 25, 35 for positive mode, and 20, 24, and 28 for negative mode. Samples were analyzed in randomized order with a technical mixture injected after every 10 samples to qualify instrument performance. Calibration was performed prior to analysis using the Pierce™ Positive and Negative Ion Calibration Solutions (Thermo Fisher Scientific). Calibration was performed prior to analysis using the Pierce™ Positive and Negative Ion Calibration Solutions (Thermo Fisher Scientific).

### 2.9 Data analysis

Acquired data was then converted from raw to mzXML file format using Mass Matrix (Cleveland, OH, USA). Discovery mode alignment, feature identification, and data filtering were performed using Compound Discoverer 3.0 (Thermo Fisher Scientific). Metabolite assignments were performed using accurate intact mass (sub-10 ppm), isotopologue distributions, and retention time/spectral comparison to an in-house standard compound library (MSMLS, IROA Technologies, NJ, USA) using MAVEN (Princeton, NJ, USA). (Melamud et al., 2010). A coefficient of variation < 20% in the technical mixtures was used as a threshold for reporting metabolites.

Metabolite molar concentrations were determined by multiplying the ratio of the monoisotope to the respective spiked in isotoplogue standard by the standard known concentration, with a subsequent correction for dilution factor during extraction as reported.[45]

Lipidomics data were analyzed using LipidSearch 5.0 (Thermo Scientific), which provides lipid identification on the basis of accurate intact mass, isotopic pattern, and fragmentation pattern to determine lipid class and acyl chain composition.

Graphs, heat maps and statistical analyses (either T-Test or ANOVA), metabolic pathway analysis, PLS-DA and hierarchical clustering was performed using the MetaboAnalyst 4.0 package. (Chong et al., 2018) XY graphs were plotted through GraphPad Prism 8 (GraphPad Software Inc., La Jolla, CA, USA). Pathway graphs were prepared on BioRender.com.

Fuzzy c-means clustering was performed using the R package ‘Mfuzz’ (v2.20.0) using 4 centers, and m value of 1.5, and a min.acore of 0.7.

Metabolite pathway enrichment analysis was performed using the Peaks To Pathways of MetaboAnalyst 5.0. Feature regions identified with the longitudinal patterns selected from C-Means Clustering were subjected to pathway analysis and enrichments were plotted a pie-charts demonstrating pathway enrichment as a function of observed vs total pathway hits.

## 3. RESULTS

### 3.1 Metabolomic changes observed in blood vary as a function of training intensity and duration

In consideration that lactate threshold and the shift between substrate and pathway utilization is dependent upon training status and performance capacity, we sought to identify circulating metabolite profiles of aerobic and anaerobic exertion in the whole blood of elite World Tour professional cyclists during a team training camp using mass spectrometry-based metabolomics. To determine lactate threshold, these cyclists first underwent a graded exercise test (GXT) to volitional exhaustion, in which progressive increases in power output on an ergometer were accompanied by whole blood lactate measurements (**Figure 1A**). A range of lactate thresholds from 3.75 to 6.5 watts per kilogram was observed in this group, demonstrating variation in metabolic capacity even at the elite level (**Figure 1B**). Guided by the lactate thresholds determined in the GXT, these cyclists subsequently completed a 180 km aerobic training session within a power output range beneath LT (**Figure 1B**). Importantly, guidance on training power output ranges would not be supported by heart rate monitoring alone, as circulating lactate only moderately correlated with heart rate over all measured power output ranges (R^2^ = 0.53, p < 0.0001), and this correlation depended on functional output (**Suppl. Figure 1**). As revealed through metabolomics, blood samples taken before and after the GXT could be distinctly grouped based on metabolite signatures using unsupervised Principal Component Analysis (PCA) (**Figure 1D**). A partial least squares discriminant analysis (PLS-DA) model was developed to identify metabolic characteristics of each time point and highlighted glycolysis (pyruvate, lactate), TCA cycle (malate, succinate), nucleotide/nicotinamide metabolism (xanthine, kynurenic acid, ADP-ribose, phosphate), oxidative stress (cystine, γ-glutamyl-alanine) and fatty acid oxidation (β-hydroxybutyrate, AC(2:0), AC(12:0), AC(16:2)) as top contributors to the metabolic character of samples isolated before and after the GXT (**Figure 1E**). In comparison, PCA was able to distinguish samples taken before and after the long aerobic training session, with a few samples clustering in the opposite group possibly indicating varying levels of exertion or recovery (**Figure 1F**). Time points from this training test were predominantly distinguished by acylcarnitines, intermediates of fatty acid oxidation (**Figure 1G**).

**Figure 1.**
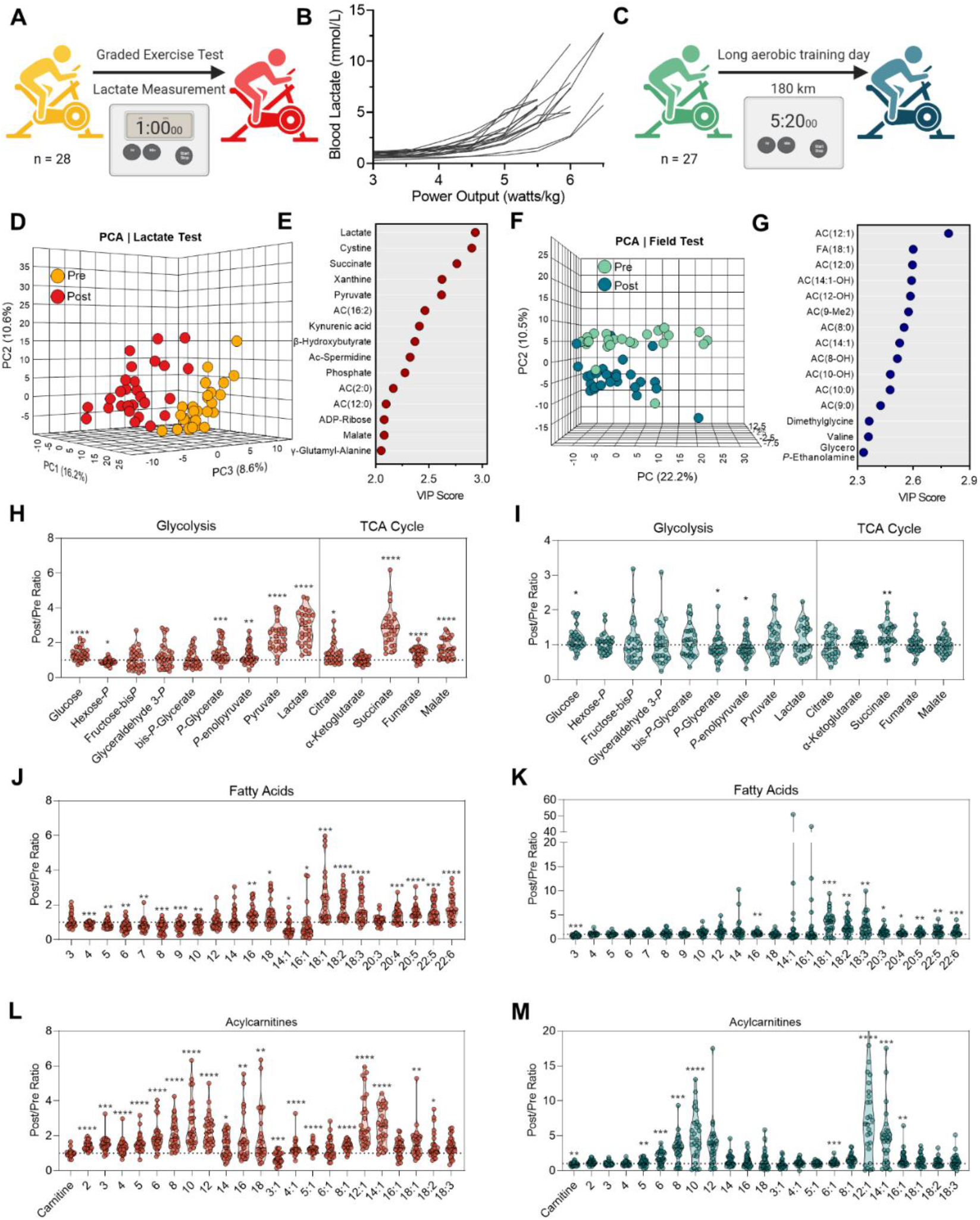
Metabolic signatures of short/high-intensity and long/low-intensity training regimens. (A) During training camp, whole blood from 28 Elite World-Tour cyclists was sampled before and after a one-hour graded exercise test on an ergometer. (B) Whole blood lactate measurements (millimolar) as a function of normalized power output (watts/kg) during the test. (C) During the same training camp, whole blood was sampled from 27 of the cyclists before and after a 180 km field test maintained in aerobic regimen beneath the lactate threshold. Multivariate analyses including Principal Component Analysis (PCA) and Variable Importance in Projection (VIP of Partial Least Squares Discriminant Analysis (PLS-DA) were performed on metabolomics data generated from the graded exercise test (D) and (E), or aerobic field test (F) and (G), respectively. Individual cyclist fold changes (Post/Pre) for metabolites involved in glycolysis and the tricarboxylic acid (TCA) cycle are shown as violin plots for the (H) graded exercise test and (I) aerobic field test. Individual cyclist fold changes (Post/Pre) for free fatty acids are shown as violin plots for the (J) graded exercise test and (K) aerobic field test. Individual cyclist fold changes (Post/Pre) for acylcarnitines are shown as violin plots for the (L) graded exercise test and (M) aerobic field test. P-values (two-tailed paired T-test) for Post/Pre comparison are indicated as *<0.05, **<0.01, ***<0.001, ****<0.0001.

In addition to the accumulation of lactate during the GXT (intra-individual fold changes of 2.91±0.94), higher levels of glucose (1.35 ± 0.38), phosphoglycerate (1.48 ± 0.59), phospoenolpyruvate (1.27 ± 0.52), and pyruvate (2.34 ± 0.78), in conjunction with lower levels of hexose phosphate (0.81 ± 0.50), indicate ongoing glycogenolysis and committal to glycolysis during this exercise test (**Figure 1H**). The accumulation of succinate (2.85 ± 1.06) during the GXT was comparable to lactate, while additional carboxylic acids citrate (1.38 ± 0.61), fumarate (1.36 ± 0.34), and malate (1.66 ± 0.54) were also increased (**Figure 1H**). These metabolic profiles were distinct from those observed in the 180 km aerobic training session, where only glucose (1.17 ± 0.32) and succinate (1.28 ± 0.37) were significantly higher after the test, though to a lower extent (**Figure 1I**).

Blood profiles of fatty acids and acylcarnitines differed between these two exercise modes as well. While long chain fatty acids (LCFA) accumulated during both exercise modes, the extent of accumulation was higher after the 180 km aerobic training session and indicated more fatty acid mobilization for energy generation (**Figures 1J** and **1K**). Increased levels of short (SCFA) and medium chain fatty acids (MCFA) after the GXT, however, suggest incomplete fatty acid oxidation during this cycling test to volitional exhaustion. In support, post-GXT blood had more abundant short chain acylcarnitines (SCAC), while blood post-180km aerobic training had no changes to SCAC and higher levels of medium chain acylcarnitines (MCAC) (**Figure 1L** and **1M**). These findings support more reliance on fatty acid oxidation during the entirety of the long aerobic training session, as further exemplified by lower levels of circulating carnitine (0.86±0.22) in comparison to the GXT (**Figure 1L** and **1M**).

### 3.2 Metabolomic changes vary based on overall progression of multistage World Tour race

We next sought to determine blood metabolic signatures in five members of this UCI elite World Tour Team during competition in a World Tour multi-stage race. To determine how these signatures change as a function of exertion both during individual stages and as a progression throughout the entire competition, whole blood samples were collected prior to and immediately upon completion of Stages 1, 4, and 6 that were characterized by distance/elevation of 143 km/60 m, 214.5 km/3,600m, and 127.5 km/4,300m, respectively. Once collected, samples were analyzed by untargeted metabolomics and lipidomics using high-throughput mass spectrometry (**Figure 2A**). Volcano plots indicated that, as the race progressed, fewer metabolites were significantly depleted during the stage (30, 26, and 11 significant features were higher before the beginning of Stages 1, 4, and 6, respectively), while Stage 6 – the most difficult stage – elicited the largest increase in circulating metabolites (**Figure 2B**). Untargeted profiles were hierarchically clustered to assess relative abundances as a function of stage and time (**Figure 2C**). From the untargeted metabolomics data, spectra for 1,971 putative metabolites were manually curated to identify 290 named compounds. PLS-DA of this sample set separated timepoints chronologically across the Component 1 axis (which explained 16.2% of the co-variance; **Figure 2D**, left). Samples obtained from all but one of the cyclists after Stages 4 and 6 begin to deviate from a linear progression and spread along the Component 2 axis. Top metabolites that influenced this clustering pattern contained primarily amino acids, free fatty acids, and acylcarnitines (**Figure 2D**, right).

**Figure 2.**
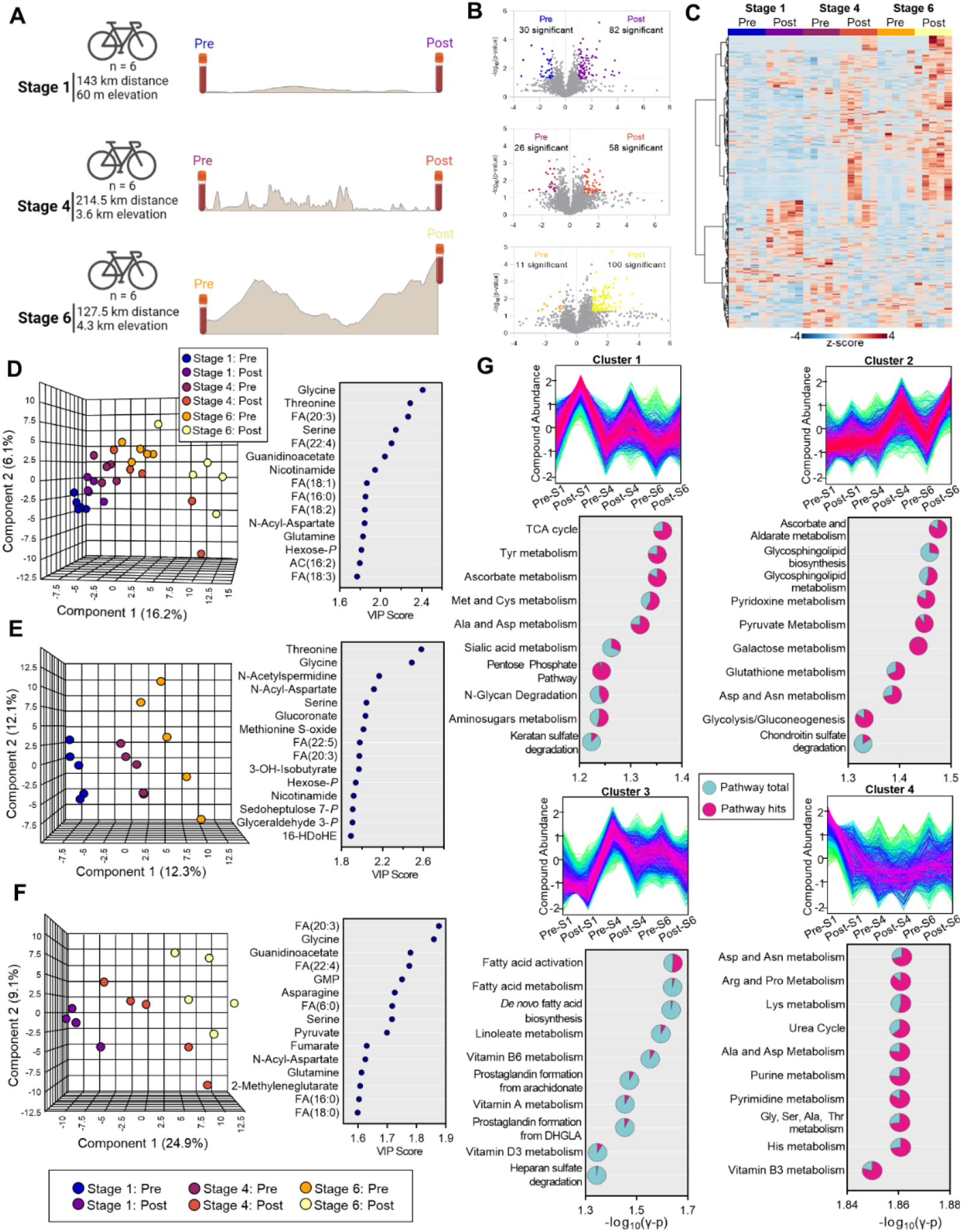
Metabolomics of a multi-stage World Tour cycling race. (A) Whole blood samples were isolated from cyclists before and after Stage 1, 4, and 6 of a consecutive 7 stage race and analyzed by mass spectrometry. (B) Volcano plots for Stage 1, 4, and 6 from top to bottom, respectively, with the number of significantly changed metabolites (fold change>2, p < 0.05 indicated in the plot. (C) Hierarchical clustering analysis of features identified by metabolomics and lipidomics. (D) Partial least squares-discriminant analysis (PLS-DA) of identified metabolites and lipids before and after each stage (left) along with the top 15 compounds by Variable Importance in Projection (VIP) (right). PLS-DA of identified metabolites and lipids (E) before and (F) after each stage (left) along with the top 15 compounds by VIP (right). (G) Four distinct longitudinal signatures were determined by fuzzy c-means clustering. Compounds with patterns matching the top 4 clusters were analyzed to determine significantly enriched pathways in each cluster. Pathways for each cluster are organized by –log10(γ p-value), with number of pathway hits (pink) plotted as a fraction of pathway total (turquoise).

Considering that the clustering pattern mirrored the chronologic progression of stages, we then analyzed individually the pre and post time points. PLS-DA separated samples taken before Stages 1, 4, and 6, indicating a progressive change in baseline blood signatures throughout the race (**Figure 2E**, left). These changes related primarily to amino acids (glycine, threonine, serine, 3-hydroxyisobutyrate), nitrogen metabolism (N-acetylspermidine), fatty acid metabolism (FA(22:5, 20:3)), glycolysis/energy (glyceraldehyde 3-phosphate, nicotinamide), and oxidative stress (methionine S-oxide, hexose phosphate, sedoheptulose 7-phosphate) (**Figure 2E**, right). Meanwhile, samples taken after each stage were even more distinguishable as the race progressed. Two cyclists sampled after Stage 4 clustered with the post-Stage 6 group, suggesting that they were accumulating markers of fatigue earlier on in the race (**Figure 2F**, left). The relative proximity of these groups depended predominantly on free fatty acids FA(20:3, 22:4, 6:0, 16:0, 18:0)), amino acids (glycine, asparagine, serine, glutamine), and energy metabolites (GMP, pyruvate, fumarate) (**Figure 2F**, right).

To identify patterns of metabolite changes throughout the race, we performed c-means clustering (**Figure 2G**). Cluster 1 contained molecules that increased during each stage, though to a lower extent as the race went on (**Figure 2G**, upper left). Pathway analysis of molecules in this cluster were enriched in energy metabolism (TCA Cycle), anabolism/oxidative stress response (Pentose Phosphate Pathway (PPP), ascorbate metabolism, methionine and cysteine metabolism), amino acid metabolism (tyrosine, methionine, cysteine, alanine, aspartate), and protein glycosylation (sialic acid and aminosugar metabolism, N-glycan and keratan sulfate degradation). Metabolites contained in cluster 2 opposed this trend, increasing primarily after the completion of Stages 4 and 6 (**Figure 2G**, upper right). This cluster pertained to oxidative stress (ascorbate metabolism, glutathione metabolism) and energy metabolism (pyridoxine and pyruvate metabolism, glycolysis and gluconeogenesis). Meanwhile, cluster 3 described molecules that are elevated after the completion of Stage 1 and progressively decrease throughout Stage 6 and related primarily to fatty acid activation (**Figure 2G**, lower left). Finally, molecules that decreased after the beginning of the race and did not predominantly recover through Stage 6 were described by cluster 4 and enriched for the metabolism of amino acids, nucleotides, nicotinamide (**Figure 2G**, lower right).

### 3.3 Longitudinal Effects of Cycling on Energy Metabolism

Considering the enrichment of glycolysis, the PPP, and the TCA cycle from the systematic analysis of untargeted metabolomics, we then manually interrogated each of these pathways (**Figure 3A**). Intermediates of the oxidative phase of the PPP including gluconolactone-6-phosphate and 6-phosphogluconate tend to increase after each stage (or significantly increase after Stage 6 or 4, respectively) (**Figure 3B**). Meanwhile, non-oxidative intermediate sedoheptulose 7-phosphate decreased, while the pool of pentose phosphate isobars tended to increase after each stage (**Figure 3B**). At steady state, these results suggest that the PPP is activated in response to cycling, which is supported by the progressive increase in glucose 6-phosphate throughout the race (**Figure 3C**). Glucose phosphorylation by hexokinase commits it towards intracellular catabolism through both the PPP and glycolysis. Significantly increased glucose after Stage 1 indicates ongoing glycogenolysis to fuel these processes. Utilization of glycolysis is especially apparent in Stage 1 given the decrease in glycolytic intermediates and increase in end-products pyruvate and lactate. (**Figure 3C**). Interestingly, the accumulation of lactate after each successive stage is lower, suggesting either a routing of carbon into the mitochondria to fuel oxidative phosphorylation, or a progressive loss in glycolytic capacity in these cyclists as fatigue accumulates. While citrate increases (significantly after Stage 6), additional TCA cycle intermediates succinate, fumarate, and malate accumulate after each stage, though to a lesser degree throughout the race (**Figure 3D**). Meanwhile, the respective amino acid products of transamination reactions (alanine/pyruvate, aspartate/malate, glutamine/glutamate/α-ketoglutarate) inversely mirror their carboxylate counterparts, suggesting that these amino acids are consumed via transamination reactions to fuel the TCA cycle and maintain nitrogen homeostasis (**Figure 3E**).

**Figure 3.**
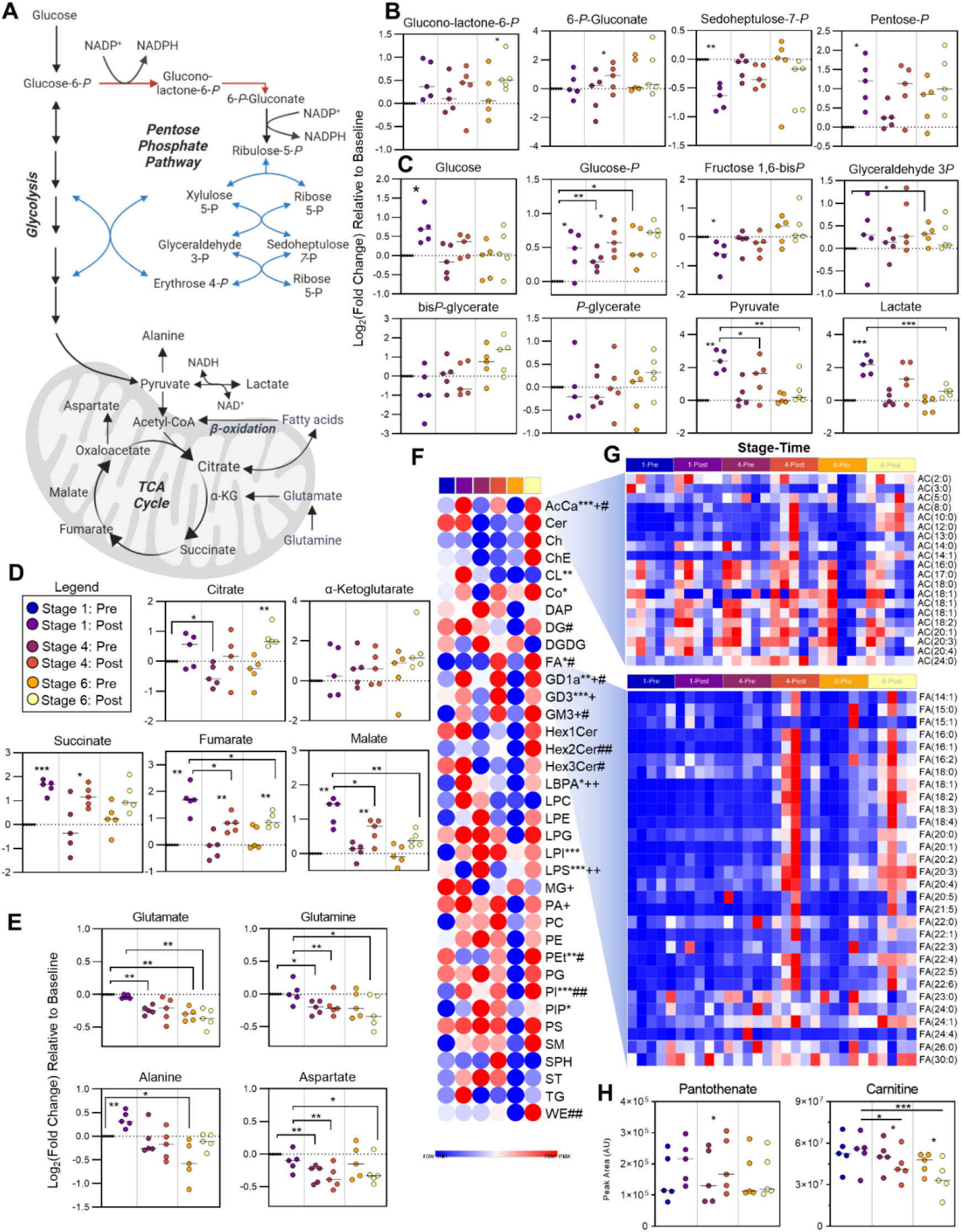
Energy metabolism. (A) A pathway overview of energy metabolism is shown, along with stage comparisons of individual metabolite levels in (B) the Pentose Phosphate Pathway (PPP, oxidative phase in red, non-oxidative phase in blue), (C) glycolysis, and the (D) Tricarboxylic Acid (TCA) Cycle and the (E) amino acid transamination products. P-values are depicted as *<0.05, **<0.01, ***<0.001. (F) A heat map for group averages of the summed total for lipid classes is shown. (G) Relative levels at each stage/time point of acylcarnitines (top) and fatty acids (bottom) for each cyclist are shown. P-values for Stage 1 Pre/Post comparison (*<0.05, **<0.01, ***<0.001), Stage 4 Pre/Post comparison (+) and Stage 6 Pre/Post comparison (#) are shown. (H) The peak areas at each stage/time point for Coenzyme A precursor pantothenate and carnitine are shown.

### 3.4 Longitudinal Effects of Cycling on Lipid Metabolism

In addition to the use of glucose and amino acid-derived carbon to fuel the TCA cycle, fat also serves as a fuel source especially under aerobic conditions of exercise. As such, we performed untargeted lipidomics to assess the relative levels of lipids throughout the race, which enabled relative quantification of 975 lipids after data curation. Of all the lipid classes profiled, AC were the only class to significantly change after each stage (**Figure 3F**). Analysis of the relative levels of individual AC revealed marked accumulations of SCAC and MCAC after Stage 6, and two cyclists finished this stage with high levels of LCAC (**Figure 3G**). The levels of FA were even more distinctly elevated after Stage 6 in all cyclists, though this accumulation begins to appear after Stage 4 (**Figure 3G**). The oxidation of FA is in part dependent in part on the availability of carnitine, which significantly decreased throughout the course of the race. Meanwhile the levels of the Coenzyme A precursor, pantothenate, remained unchanged aside from a slight, though significant, increase after Stage 4 (**Figure 3H**).

### 3.5 Individual Cyclist Metabolomics as a Function of Performance

The penultimate stage of the race consisted of a 127.5 km course through a mountainous region that finished with a first category climb (7% average grade) over the final 7.5 km (**Figure 4A**). Cyclists 1 and 2 managed to outpace the other 3 throughout this stage, which was driven in part by the final vertical ascent (**Figure 4B**). To identify metabolite signatures that associate with performance during this period of the race, we correlated average speed with metabolite levels in blood sampled immediately after Stage 6. Top correlates pertained predominantly to oxidation of fatty acids (AC(18:2), AC(18:1), AC(16:0), AC(18:0), AC(18:3), AC(3:1), FA(6:0), and pantothenate), nitrogen homeostasis (5-hydoxyisourate, 5-methylthioadenosine, 4-acetoamidobutanoate, lysine, spermidine, and spermine), and energy metabolism (glyceraldehyde 3-phopshate, succinate) (**Figure 4C** and **4D**). To expand analysis of personalized metabolic profiles across the entirety of the race, sparse Partial Least Squares Discriminant Analysis (sPLS-DA) revealed a similar trend as the speed and output data, with the fastest cyclists 1 and 2 clustering together along the Component 2 axis (**Figure 4E**). The top 10 metabolites that contribute to the clustering pattern along this axis were enriched for intermediates of fatty acid oxidation (**Figure 4F**). Indeed, Cyclists 1 and 2 maintained the highest levels of CoA-precursor pantothenate and LCAC throughout the duration of Stages 1, 4, and 6 (**Figure 4G**). Cyclist 1, who was the top performing cyclist in this cohort, had persistently high levels of carnitine and the lowest levels of MCAC and LCFA (**Figure 4G** and **4H**), suggesting maintenance of mitochondrial capacity and fatty acid oxidation throughout the race. In support, this cyclist also finished Stage 6 with the lowest levels of glycolytic intermediates, lactate, and succinate (**Figure 4I** and **4J**), along with a larger pool of NAD(H) (**Figure 4K**), indicating a lower fatigue status that could sustain a faster pace towards the end of the race. This same cyclist also displayed the highest levels of arginine and polyamines (spermine – **Suppl. Figure 2.A**), argininosuccinate, 5-methylthioadenosine (**Suppl. Figure 1.B**), and second highest levels of citrulline, but lowest levels of S-Adenosyl-methionine, creatinine and phosphocreatine (**Suppl. Figure 2.B**) across all other team members.

**Figure 4.**
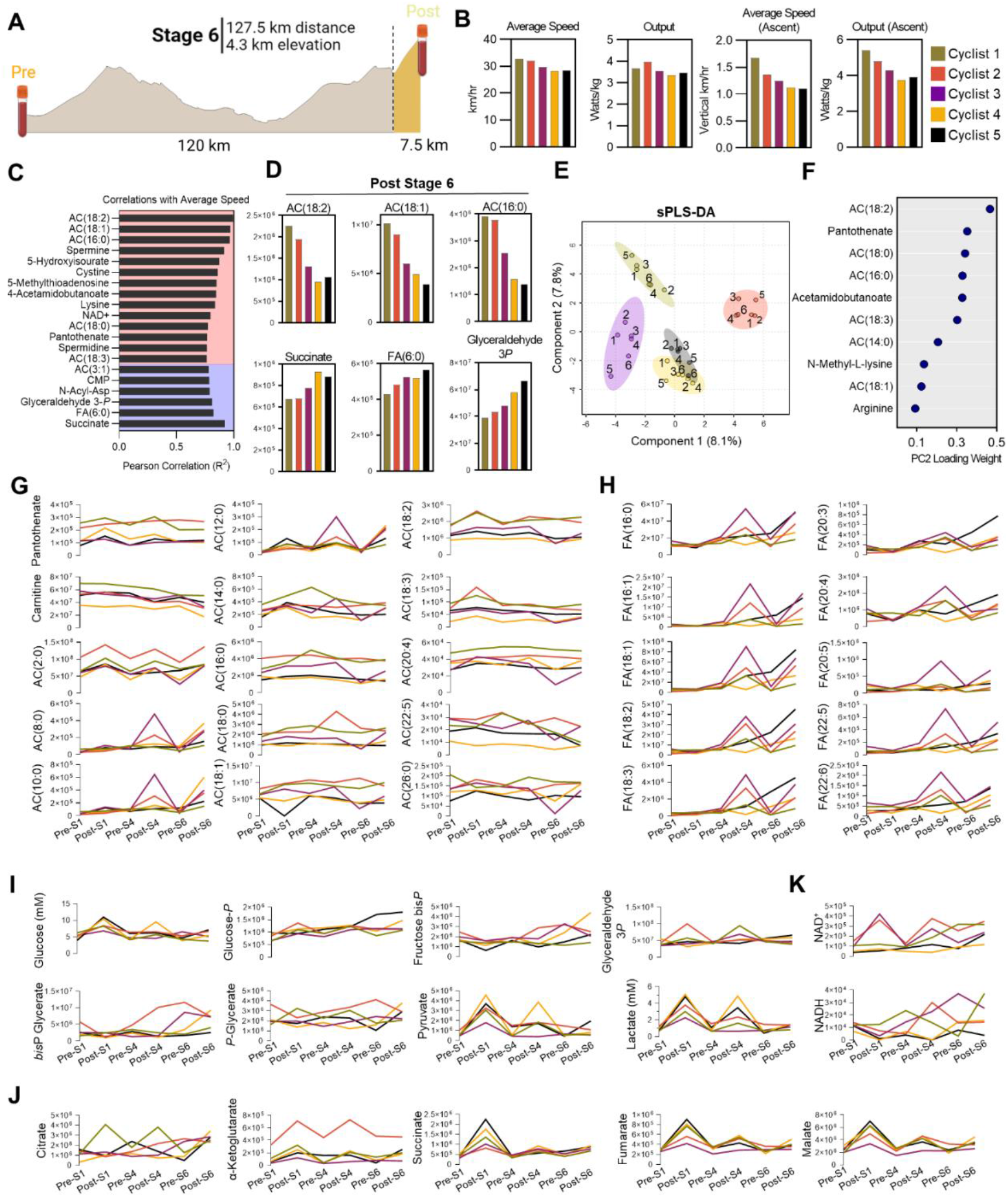
Individual Cyclist Analysis. (A) Metabolite, lipid, and Training Peaks functional cycling data for the entirety of Stage 6, along with Training Peaks during the final 7.5 km climb were analyzed. (B) Average speed and output for Stage 6, as well as the average speed and output of the final 7.5km are plotted by individual cyclist. (C) Pearson Correlation coefficients (R^2^) for the top 20 Post-Stage 6 metabolite correlates with average speed during Stage 6 are shown, with positive correlates indicated in red background and negative correlates indicated in blue background. (D) Abundances (y-axis values are peak area top, given in arbitrary units) for each cyclist of the top 3 positive (top) and negative (bottom) correlates with average speed are shown. (E) Sparse Partial Least Squares Discriminant Analysis (sPLS-DA) is shown, with samples color coded according to cyclist and time point indicated as 1-6, where 1 = Stage 1-Pre consecutively up to 6 = Stage 6-Post. (F) The top 10 loadings for principal component 2 are plotted. Longitudinal profiles during the course of the race for (G) acylcarnitines, (H) free fatty acids, (I) glycolysis, (J) TCA cycle, and (K) NAD+/NADH in each cyclist are shown as line graphs, with time points indicated on the bottom x-axis and abundance indicated on the y-axis as peak area (arbitrary units) except for glucose and lactate, for which the absolute concentrations are reported (millimolar).

Most notably, Cyclist 1 showed the highest levels of reduced and oxidized glutathione (total glutathione pools), and methionine (**Suppl. Figure 2.C**), suggestive of the highest antioxidant capacity throughout competition among all cyclists monitored in this study.

## 4. DISCUSSION

Here we used mass spectrometry-based metabolomics to define whole blood molecular profiles associated with sustained low to medium intensity cycling during a 180 km aerobic training ride in comparison with a graded exercise test (GXT) to volitional exhaustion. To facilitate field sampling, we circumvented the needs for traditional phlebotomy and maintaining frozen samples as is traditionally required for metabolomics analyses through the implementation of dried blood sampling using volumetric absorptive microsampling[20]. Because these samples were taken from the same elite professional cyclists during a weeklong training camp prior to the season, they enable a paired comparison of molecular profiles as a function of exertion and serve to define blood profiles of humans performing at optimal capacity. Metabolite profiles of the GXT demonstrated characteristic accumulations of circulating lactate, a biomarker of performance capacity that has been traditionally used to guide training exercise [19,21]. In addition, we were also able to quantify the extent to which upstream glycolytic intermediates are modulated to sustain lactate production. The accumulation of lactate occurs at a power output when the rate of lactate production exceeds that of oxidation, termed the “lactate threshold,” which is dependent upon the ability to oxidize lactate into pyruvate for subsequent metabolism in the mitochondria[10,22]. Exercise intensity dictates demand for ATP and drives skeletal muscle metabolic responses. At high exercise intensities, glycolysis is the primary source of ATP, which is produced at a faster rate through the activity of glycolytic enzymes phosphoglycerate kinase and pyruvate kinase in comparison to mitochondrial electron transport chain-fueled ATP synthase. The latter route for ATP synthesis is dependent on mitochondrial capacity, which becomes limiting at higher workloads. Under hypoxic conditions or at high bioenergetic demand, mitochondrial respiration becomes uncoupled leading to an accumulation of succinate, which is released into the extracellular environment[7,23] as a function of intracellular proton accumulation[24]. In line with previous findings[19], we observed increases of succinate that were comparable to that of lactate during the GXT. During the 180 km training session however, when cyclists are functioning in aerobic conditions and primarily relying on fatty acid oxidation, succinate shows only modest increases while lactate remained unchanged.

High exercise intensities are predominantly sustained by carbohydrate oxidation, while low and medium intensities rely more upon catabolism of fatty acids for energy generation. Accordingly, we observed a substantially larger accumulation of LCFA including the most abundant oleic and linoleic acid [25] after the 180 km aerobic training session in comparison with the GXT. The relatively higher levels of these fatty acids after the long training session indicate a longer period of fatty acid mobilization during this training test, which are subsequently converted intracellularly between acyl-CoA and acylcarnitine species for fatty acid oxidation within the mitochondria. The inability of mitochondria to continue oxidizing fatty acids, either due to lack of oxygen availability or cessation of exercise, results in release of incompletely oxidized MCAC back into circulation. SCAC and MCAC notably accumulated significantly after the GXT in comparison to the lower intensity training session, indicating an abrupt shift in metabolism due to progressively increasing exercise intensity. The resulting accumulation of lactate exerts endocrine and autocrine actions by decreasing lipolysis[26] and mitochondrial fatty acid transport through decreased carnitine palmitoyltransferase I and II (CPT I and II) function[27]. Of note, MCAC are higher at baseline in patients with Type 2 diabetes[28,29], sepsis[30,31], and post-acute sequelae of SARS-CoV-2 infection (PASC) [32], and indicate that metabolomic signatures of acute exercise-induced fatigue resemble those of chronic diseases in which metabolic and mitochondrial dysfunction or impairment play an important pathogenic role.

When translated into a World Tour cycling competition, the metabolomic signatures presented herein enable assessment of workload and performance. For instance, the most significantly enriched pathway in Cluster 1, which described compounds that accumulate the most after Stage 1, was the TCA Cycle. This stage was characterized by long stretches of flat terrain with few hills that finished with a field sprint, which demanded higher workloads in line with the profiles seen in the GXT to volitional exhaustion. Notably, lactate and succinate both accumulated in circulation largely this stage, thus indicating the high degree of exertion and fatigue in these cyclists after the sprint. On the other hand, Stages 4 and 6 involved more climbing, and Stage 6 finished at the highest elevation of the race. Accordingly, we observed profiles indicating a predominant reliance on fat oxidation, as reflected by accumulation of fatty acids and MCAC after the stage.

While this study only profiled the longitudinal patterns of 5 professional cyclists, correlation of metabolic profiles with speed revealed interesting patterns. The slowest 2 cyclists during Stage 6 finished with the lowest levels of LCAC and the highest levels of circulating succinate, the latter of which was a marker of fatigue in the GXT. It is interesting to note that these two cyclists also had the highest levels of circulating glucose after Stage 1, as well as the highest lactate after Stages 1 and 4. This signature indicates that these two cyclists were more reliant on glycolysis early in the race, accessing glycogen storage for sustained output. As such, they may have depleted stores and began to fatigue more quickly than their teammates, thus resulting in poorer performance towards the end of the race. Meanwhile, the fastest 2 cyclists had lower succinate and higher LCAC after Stage 6, indicating ongoing aerobic fatty acid oxidation despite generating the highest power output and speed on the final ascent. Longitudinal characterization of all three sampled stages during the race revealed that these two cyclists maintained the highest levels of LCAC across this cohort, along with the highest level of CoA precursor pantothenate (Vitamin B5). Furthermore, Cyclist 1 had the lowest markers of incompletely oxidized fatty acids, along with the highest level of carnitine. This cyclist maintained the highest lactate threshold measured during training camp (**Figure 1B**) and was the lone individual who did not deviate from the metabolic trajectory in the World-Tour identified by PLS-DA (**Figure 2D**). This cyclist also had higher levels of branched-chain amino acids (BCAA) than the other cyclists, especially prominent after Stage 6 when isoleucine, leucine, and valine reached concentrations of 120, 219, and 264 µM, respectively (**Suppl. Fig 3**). In addition to its role in fatty acid oxidation, CoA plays a predominant role in BCAA catabolism, and its biosynthesis may be tied to lactate threshold. While plasma levels of BCAA have been reported to respond to exercise duration and work load [33–35], BCAA oxidation propensity is also tied to endurance capacity. [36] These results indicate that high performing elite cyclists have an increased capacity to maintain mitochondrial metabolism and oxidative phosphorylation at high workloads and substantiate prior findings that cyclists’ performance correlates with higher lactate clearance capacity at comparable power output. [12]

Sports performance has both genetic and environmental influencing factors. Indeed, elite performance does have a basis in intensive training regimens. [37] While genetic factors are not as capable of distinguishing elite athletes on their own, combination of GWAS with metabolomics using metabolic quantitative trait loci (mQTLs) analysis revealed significant associations between metabolite levels and genetic features [38]. One such mQTL is an association between the endocannabinoid linoleoyl ethanolamide and vascular non-inflammatory molecule 1 (VNN1), which functions as a panetheinase that plays an integral role in recycling of pantothenate to promote mitochondrial activity[39]. It is therefore interesting to consider the role VNN1, or additional effectors of CoA/carnitine biosynthesis and fatty acid oxidation, may play in training, fatigue, and performance, especially in light of the putative association between this pathway and cyclist performance presented here. Ultimately, the complementarity of this approach holds future promise for personalization of training regiments for sports performance and exercise prescription to treat metabolic syndrome[40] and improve cancer survivorship[41,42]. The use of dried blood sampling enables these studies and allows for applicability to the general population at large.

Although limited by its observational nature, these studies provide a unique view into metabolism of elite athletes performing at the best of their abilities both during training and in competition. Many of the cyclists profiled here are globally competitive and have stage and World Tour wins to their names. As such, these profiles provide a unique view into bioenergetics and metabolic physiology of optimal human performance.

## 5. Conclusion

The study shows that field sample collection for metabolomics analyses in professional elite athletes is feasible and informative of athlete condition and ultimate performance. The combination of lower cost high-throughput omics strategies and field sampling for remote collection without a phlebotomists can democratize omics technologies by making them accessible logistically and economically to all athletes, from recreational to elite ones. Although limited by its observational nature, these studies provide a unique view into metabolism of elite athletes performing at the best of their abilities both during training and in competition. Many of the cyclists profiled here are globally competitive and have stage and race wins to their names. As such, these profiles provide a view into bioenergetics and metabolic physiology of optimal human performance. The use of dried blood sampling enables these studies and allows for applicability to the general population at large.

## Supplementary Information

The online version contains supplementary material available at *https://doi.org//to-be-added-upon-acceptance*

## Acknowledgments

Research reported in this publication was funded by the National Institute of General and Medical Sciences (RM1GM131968 to ADA), and R01HL146442 (ADA), R01HL149714 (ADA), R01HL148151 (ADA), R01HL161004 (ADA), and R21HL150032 (ADA) from the National Heart, Lung, and Blood Institute. The content is solely the responsibility of the authors and does not necessarily represent the official views of the National Institutes of Health.

## Authors’ contribution

ISM, TN, and AD designed the studies. JLM, ISM performed exercise tests and collected the samples in laboratory and field settings. TN, FC, DS, KCH, AD performed omics analyses; TN performed data analyses and prepared figures. TN wrote the first version of the manuscript, which was finalized with AD and reviewed and approved by all co-authors.

