## Supplementary figures and images for "Metabolic Signatures of Performance in Elite World Tour Professional Cyclists"

### Supplemental Figure 1

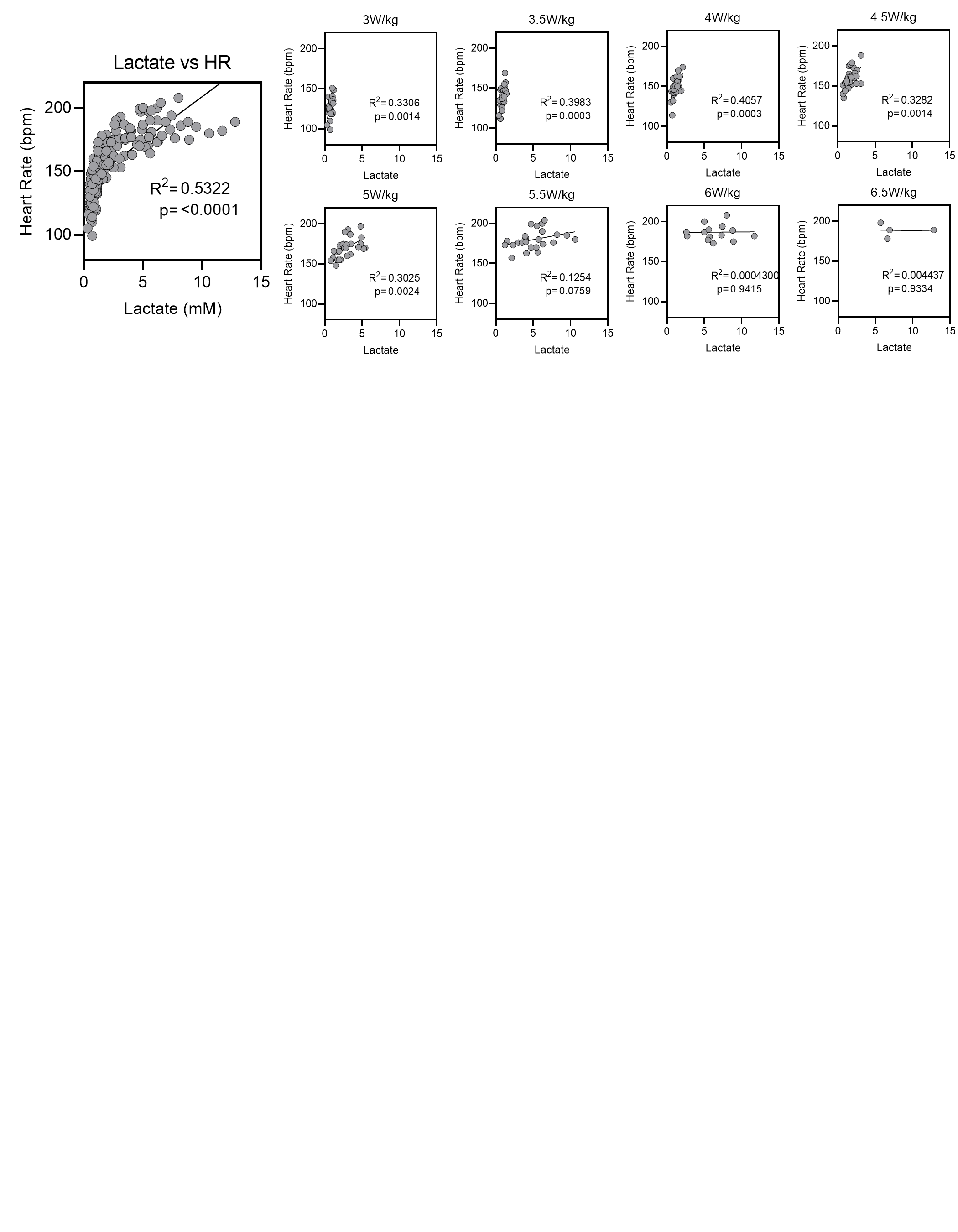

### Supplemental Figure 2

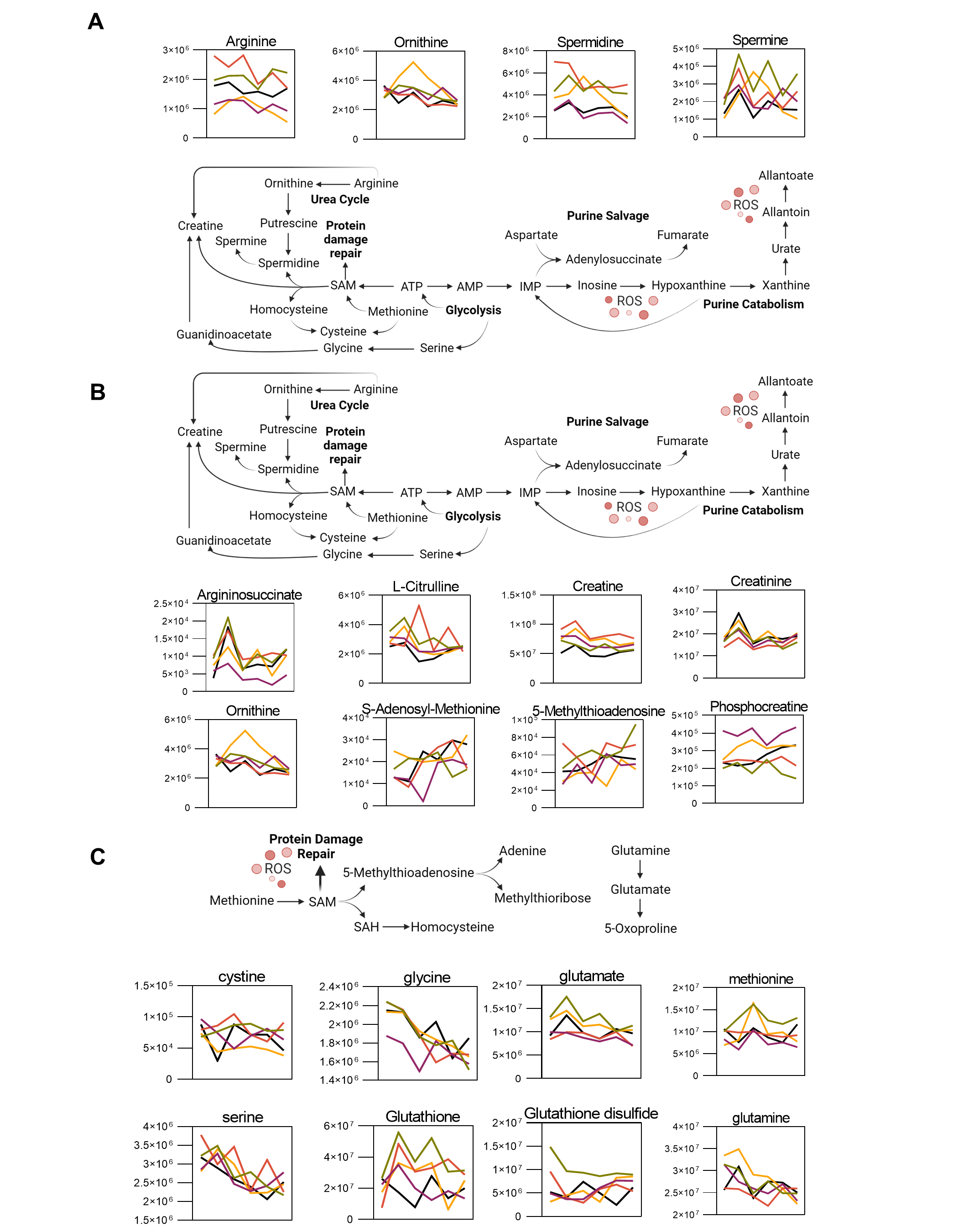

### Supplemental Figure 3

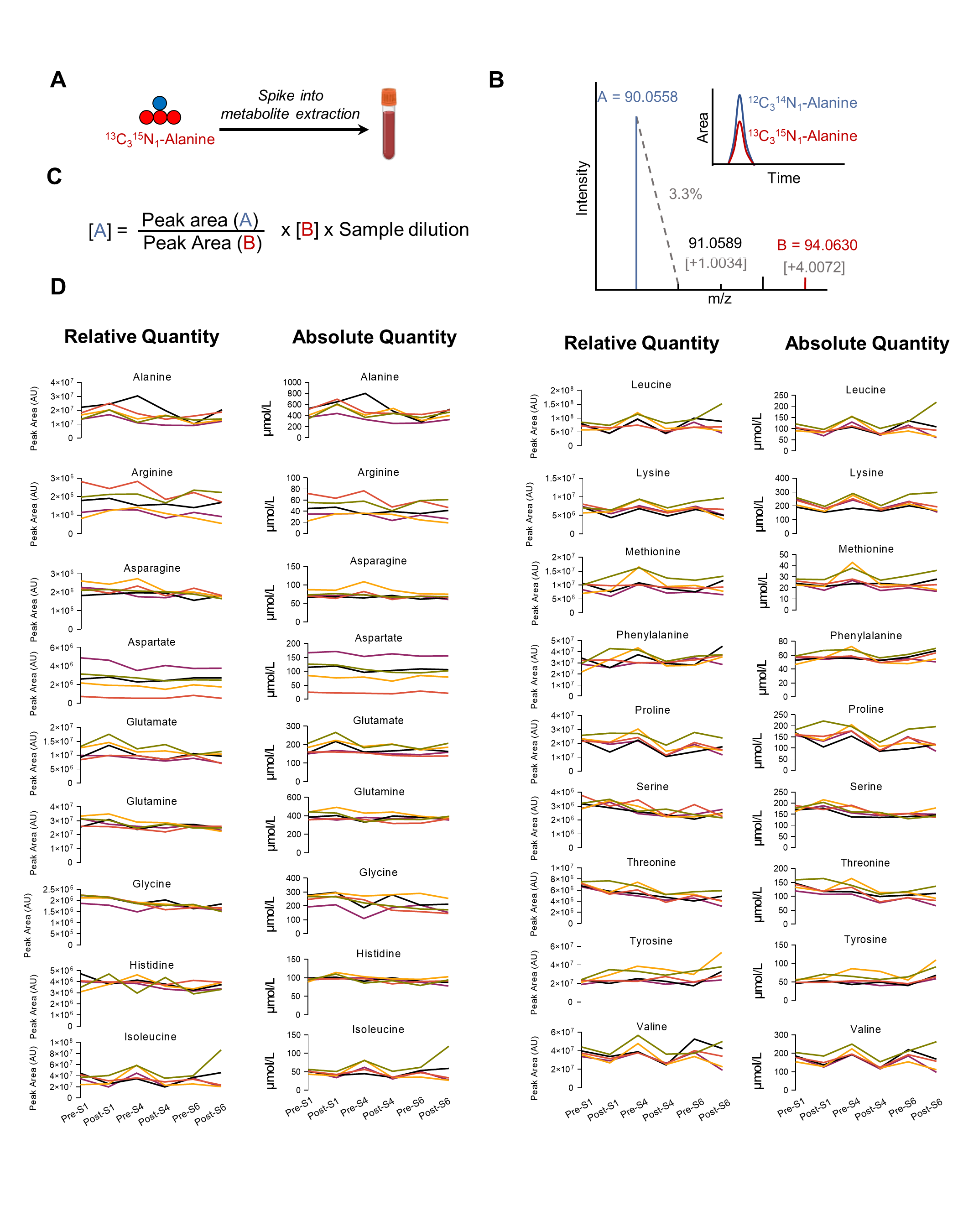
